# MUSE enables cross-species multi-omics integration that incorporates transcriptional regulatory modules

**DOI:** 10.64898/2026.05.04.722812

**Authors:** Fuka Nakae, Shintaro Yuki, Zhenan Liu, Chikara Mizukoshi, Shuto Hayashi, Teppei Shimamura, Hiroshi Yadohisa

## Abstract

Recent advances in evolutionary biology and biomedical research have promoted comparative analyses of cellular states and developmental processes across species, leading to the development of numerous cross-species alignment methods based on scRNA-seq data. However, alignments relying solely on RNA expression are strongly driven by lineage signals and cell typespecific transcriptional programs. As a result, they are limited in their ability to identify conserved regulatory modules across species and to compare regulatory logic beyond developmental lineages, thereby constraining biological interpretability.

Meanwhile, recent technological developments have enabled the acquisition of multi-omics data, including chromatin accessibility, making it increasingly feasible to analyze and interpret conservation at the level of regulatory modules across species. Nevertheless, computational methods that can integratively handle such heterogeneous omics data and enable cross-species comparative analysis in a unified framework remain insufficiently established.

To address this challenge, we propose Multi-omics Unified embedding across Species (MUSE), a novel framework for integrating multi-omics data across species. MUSE constructs a graph that captures relationships among features both within and across species, and learns a shared latent space based on this graph structure. By leveraging this integrated graph-based representation, MUSE enables cross-species alignment that preserves species-specific characteristics while reflecting similarities not only at the level of gene expression and chromatin states but also at the level of regulatory modules.

## Introduction

There has been increasing interest in cross-species analysis of single-cell RNA sequencing (scRNA-seq) data have enabled comparative investigations across species. From an evolutionary biology perspective, such analyses facilitate the identification of both conserved and species-specific features arising during evolutionary processes [1]. From a biomedical and clinical perspective, they provide a framework for inferring phenomena that are difficult to directly examine in humans due to ethical and cost constraints, such as drug responses, by leveraging experimental results obtained from model organisms [2]. Motivated by these advantages, methods for crossspecies analysis of single-omics data continue to be actively developed [3–5].

Meanwhile, recent advances in data acquisition technologies have enabled the analysis of multi-omics data that incorporate regulatory information, such as chromatin accessibility, in addition to gene expression. In particular, chromatin accessibility obtained from single-cell assay for transposase-accessible chromatin sequencing (scATAC-seq) reflects upstream regulatory states of transcription, including transcription factor binding potential and enhancer activity, and provides complementary information to scRNA-seq, which captures the outcomes of gene expression [6, 7]. Therefore, integrating scRNA-seq and scATAC-seq is expected to enable a more accurate characterization of regulatory logic, such as transcription factor activity and regulatory modules, at the single-cell level.

From an evolutionary perspective, interspecies diversity has been suggested to arise from changes in transcription factor–mediated regulatory modules [8]. Thus, the use of multiomics data including ATAC enables the analysis of conservation and divergence at the level of regulatory modules. Indeed, while cross-species alignment based on scRNA-seq primarily captures similarities in gene expression programs associated with cell types and developmental lineages [9], it has been pointed out that such approaches fail to adequately capture regulatory module-level conservation when regulatory programs have undergone evolutionary changes [8]. Furthermore, analyses based on single-cell multiome data that simultaneously measure RNA expression and chromatin accessibility have reported cases in which regulatory modules defined by chromatin accessibility and enhancer codes are highly conserved across species, even when gene expression patterns differ substantially [10]. These findings highlight the limitations of single-omics-based analyses.

Although multi-omics data enable more comprehensive analyses [11], several challenges remain in their integration. First, technical heterogeneity arises from differences in measurement technologies, data formats, and annotation standards [12]. Second, there is the problem of limited availability of comparable datasets that can be aligned across modalities [12].

To address these challenges, numerous multi-omics integration methods have been proposed to align modalities that are not measured in the same cells within a shared latent space [13–19]. Among them, graph-linked unified embedding (GLUE) is a method that incorporates prior knowledge of regulatory relationships as a graph to explicitly model correspondences between modalities. GLUE has been reported to outperform existing methods in terms of both integration accuracy and biological interpretability [20]. Such a framework, which represents inter-modality relationships as a graph and incorporates them into integration, is also promising as a foundation for cross-species data integration. However, cross-species integration presents additional challenges beyond those described above. Specifically, datasets that are comparable and properly aligned across species are even more limited, and it is necessary to integrate conserved regulatory modules while accounting for species-associated biological differences, such as evolutionary changes in gene function and regulatory relationships, as well as the presence of species-specific genes.

On the other hand, GLUE is designed under the assumption of a single species and cannot explicitly address these challenges specific to cross-species integration, making it difficult to directly apply to such settings. To overcome this limitation, we propose a novel framework termed Multi-omics Unified embedding across Species (MUSE). This method constructs a cross-species graph by introducing inter-species edges based on orthologous relationships and protein sequence similarity, while preserving the regulatory graph structure within each species. Based on this integrated graph, MUSE performs data integration that enables biologically meaningful cross-species comparisons at both the omics level and the regulatory module level.

We applied the proposed method to multi-omics data derived from the developing cerebral cortex of human, macaque, and mouse[21][22]. The results show that MUSE effectively removes species-specific batch effects while accurately preserving cell-type structure and biological characteristics. Furthermore, the method is capable of capturing both species-specific features and conserved features across species. In addition, by integrating cross-species alignment with SAMap and regulatory module analysis with SCENIC+, we show that our approach can identify conservation at the level of regulatory modules, which cannot be fully captured by conventional methods based solely on RNA expression Taken together, these results demonstrate that MUSE is a novel framework for cross-species alignment of cellular states and comparative analysis of regulatory mechanisms, enabling the integrated interpretation of both evolutionary conservation and species-specific divergence.

## Results

### Overview of MUSE

An overview of the analysis workflow using MUSE is shown in Fig. 1. We applied MUSE to multi-omics datasets consisting of scRNA/snRNA and scATAC/snATAC data derived from the developing cerebral cortex of human (mid-gestation), macaque (gestational days 80–93), and mouse (embryonic day 14.5) [21, 22] (Fig. 1). Details of data preprocessing are provided in the Methods. In the proposed MUSE framework, we construct a crossspecies guidance graph to represent within-species regulatory structure and cross-species correspondence (Fig. 1A). The guidance graph is a weighted graph that encodes relationships between features from different omics modalities based on prior knowledge. In this study, genes (RNA) and chromatin features (ATAC peaks or highly variable peaks) are treated as nodes, and edges are assigned based on known gene-regulatory region associations. Guidance graphs are first constructed independently for each species and then integrated using amino acid sequence embedding similarities derived from the protein language model Evolutionary Scale Modeling 2 (ESM2)[23], together with ortholog information (see Methods Section 4.2, Cross-species guidance graph, for details). This procedure preserves intra-species regulatory graph structures while enabling the construction of a crossspecies guidance graph that reflects conserved regulatory architecture across species.

**Fig. 1.**
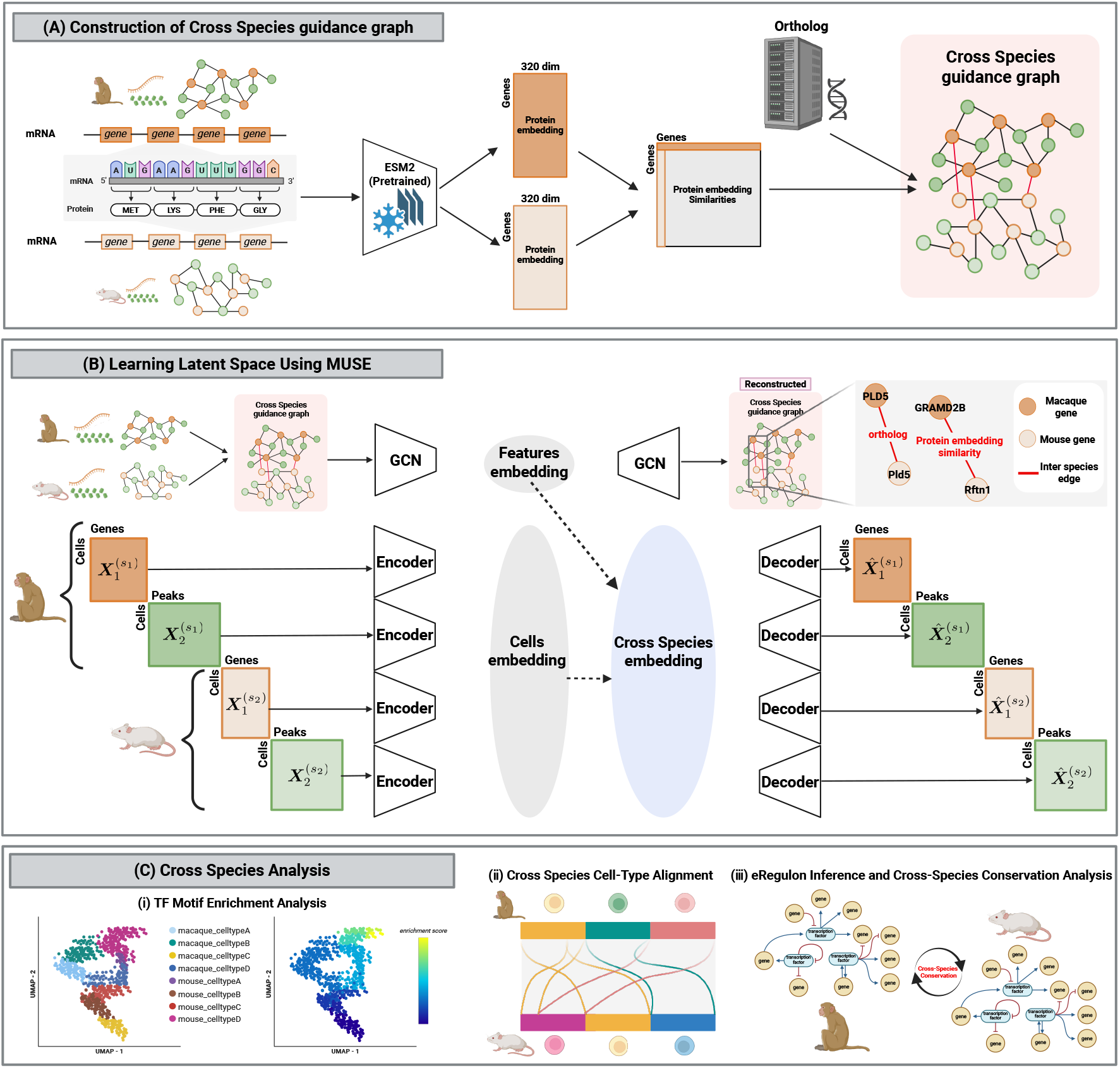
Overview of the MUSE framework for cross-species multi-omics integration. (A) Cross-species edges are introduced based on orthology and similarity between protein sequence representations, forming a cross-species guidance graph that preserves species-specific regulatory structures while enabling inter-species correspondence.(B) The model jointly learns a shared latent representation through graph-constrained variational embedding, removing species-driven batch effects while maintaining biological structure. (C) The learned latent space enables downstream analyses, including cross-species cell-type alignment, conserved and divergent regulatory module identification, and regulatory network comparison.

Using the constructed cross-species guidance graph together with multi-omics observations as input, MUSE learns a shared latent space across species (Fig. 1B). MUSE jointly models cell-level latent representations inferred from data and feature-level representations constrained by the guidance graph (genes and peaks), thereby producing aligned embeddings across both modalities and species. This framework simultaneously reduces species-specific batch effects and highlights conserved regulatory programs, yielding lowdimensional representations suitable for downstream analyses (Fig. 1C).

Using the learned latent space, we performed cross-species analyses. First, the latent representations of each cell learned by MUSE were visualized using UMAP. The cells were then colored according to their motif enrichment scores. By examining the distribution of these scores along the latent axes, we identified and interpreted the biological processes captured by the latent representations. Second, to quantify cell type concordance across species, we computed cross-species alignment scores and evaluated the preservation of corresponding relationships. Third, we interpreted the results by integrating the regulatory modules extracted by SCENIC+ with the cross-species alignment results.

### Performance Comparison Between the Proposed Method and Baselines

To evaluate the impact of introducing a cross-species guidance graph on integration performance and the preservation of biological structure, we compared MUSE with two baselines (Baseline 1 and Baseline 2) using data from two species (Fig. 2). The dataset consisted of cerebral cortex data from macaque and mouse [21].

**Fig. 2.**
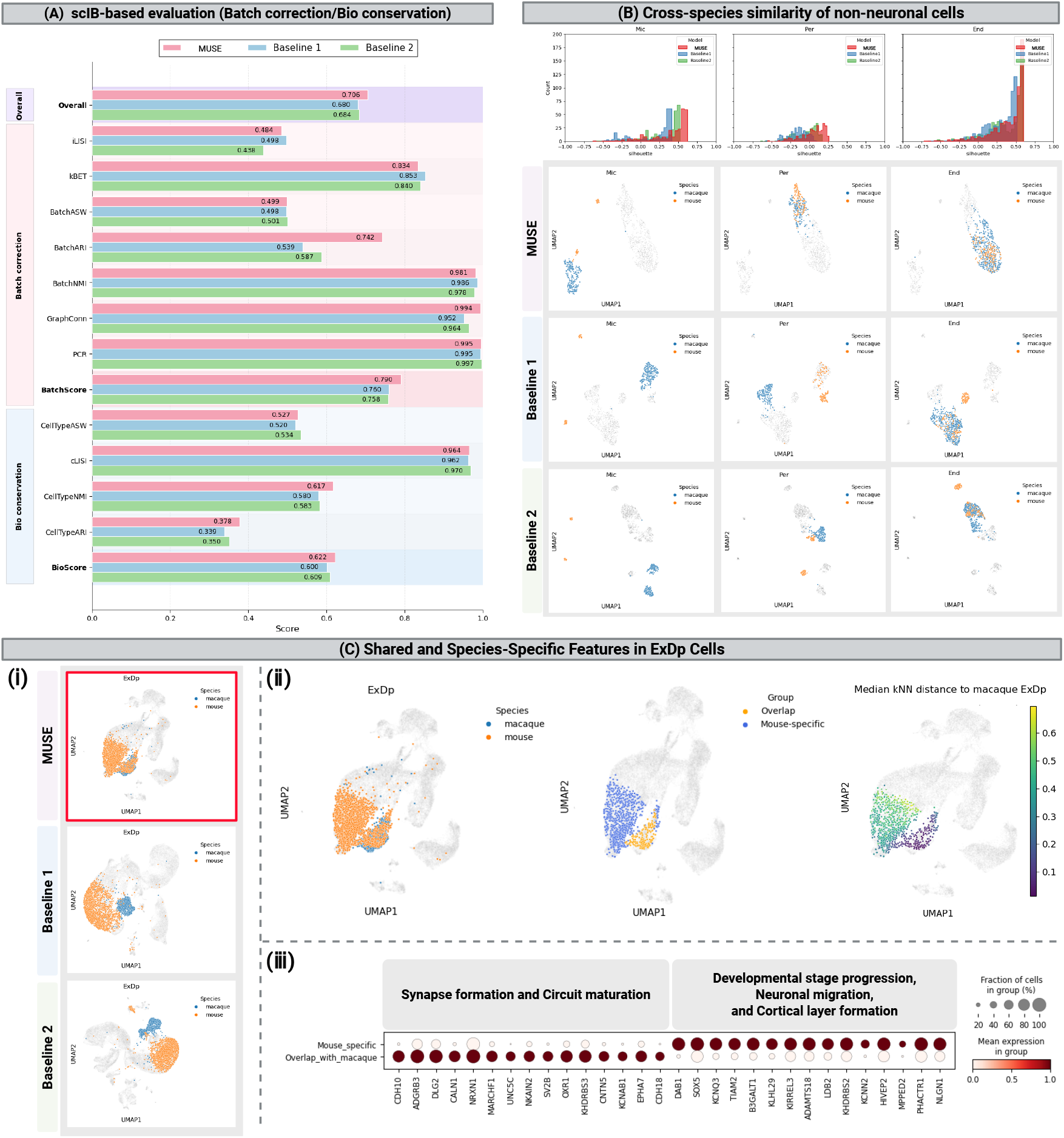
Benchmark results. (A) Quantitative evaluation using scIB metrics. MUSE achieves the highest overall performance, indicating improved batch correction and preservation of biological structure. (B) Silhouette width distributions for non-neuronal cell types (e.g., microglia, pericytes, and endothelial cells), showing improved crossspecies alignment with MUSE.UMAP visualization of non-neuronal cell types colored by species labels. MUSE aligns cells of the same type across species, whereas baseline methods show species-driven separation. (C) Differential expression analysis of mouse ExDp cells classified into “Overlap with macaque” and “Mouse specific” groups based on distances in the MUSE latent space.

An overview of the baseline methods is shown in Fig. S1. The key difference between the proposed method and the two baselines lies in the structure of the guidance graph incorporated into the model. In MUSE, guidance graphs constructed for each species are integrated based on amino acid sequence embedding similarity derived from ESM2 and ortholog information. In Baseline 1, the guidance graphs for each species remain independent. In Baseline 2, the guidance graphs are integrated using randomly assigned cross-species edges.

We first evaluated whether the proposed method can remove species-driven batch effects while preserving cell-type structure, using scIB metrics (Fig. 2A). In the overall score, MUSE achieved the highest average score, outperforming both baselines (MUSE: 0.706; Baseline 1: 0.680; Baseline 2: 0.684; Fig. 2A). From the perspective of batch correction, MUSE also achieved the highest BatchScore (MUSE: 0.790; Baseline 1: 0.760; Baseline 2: 0.758; Fig. 2A). In terms of biological conservation, as quantified by BioScore, MUSE again showed the highest performance (MUSE: 0.622; Baseline 1: 0.600; Baseline 2: 0.609; Fig. 2A). These results indicate that the proposed method improved species mixing while preserving biological structure.

Next, to assess whether cross-species similarity is captured at the level of cell types, we computed silhouette widths for non-neuronal cell types and visualized their distributions (Fig. 2B). Higher silhouette values indicate that cells are closer to cells of the same type and well separated from other types. Non-neuronal cell types (e.g., microglia, pericytes, and endothelial cells) are known to exhibit stronger conservation across species compared to neuronal populations [24], and thus their alignment in the latent space is expected.

Consistent with this expectation, the distribution of silhouette scores for MUSE was shifted toward higher values compared to the baselines across all examined cell types. We further visualized these cell types using UMAP, colored by species (Fig. 2B). Rows correspond to methods (MUSE, Baseline 1, Baseline 2), and columns correspond to non-neuronal cell types. In MUSE, cells of the same type from different species were located in close proximity, indicating successful crossspecies alignment. In particular, pericytes (Per) and endothelial cells (End) showed nearly complete overlap between species. In contrast, Baseline 1 exhibited separation between species even within the same cell type, while Baseline 2 often showed fragmentation even within the same species and cell type.

We further focused on ExDp cells and visualized them using UMAP colored by species (Fig. 2C). In the baseline methods, macaque and mouse cells were clearly separated, whereas in MUSE, a subset of mouse cells overlapped with macaque cells. Based on this observation, we performed differential expression analysis between two groups of mouse ExDp cells: those overlapping with macaque cells in the latent space (“Overlap with macaque”) and those that did not (“Mouse specific”).

Specifically, based on distances in the MUSE latent space, we defined the typical range of pairwise distances among macaque ExDp cells and used this as a reference. Mouse ExDp cells whose distances to macaque ExDp cells fell within this range were defined as “Overlap with macaque”, while the others were defined as “Mouse specific” (Fig. 2C(ii)).

In the Overlap with macaque group, genes such as *NRXN1, CDH10, CDH18*, and *EPHA7* were highly expressed (Fig. 2C(iii)). These genes have been reported to be involved in functional synapse formation, cell-cell adhesion, and neural circuit assembly [25–27]. This suggests that the Overlap group represents ExDp cells in a stage associated with synaptic maturation and circuit formation. Core transcriptional programs related to synapse formation are known to be relatively conserved across mammals [24], which may explain the alignment of these cells with macaque ExDp cells in the latent space.

In contrast, the Mouse specific group showed high expression of genes such as *DAB1, SOX5, TIAM2*, and *ADAMTS18* (Fig. 2C(iii)). *DAB1* is a key component of the Reelin signaling pathway essential for cortical layer formation, and *SOX5* is a transcription factor involved in the differentiation of deep-layer cortical neurons [28, 29]. *TIAM2* and *ADAMTS18* are associated with cell migration and extracellular matrix remodeling [30, 31], suggesting that this group reflects a developmental or structural remodeling state.

These results suggest that the Mouse specific group is associated with transcriptional programs related to developmental processes, rather than mature, conserved ExDp states. The observed separation between mouse and macaque cells is consistent with previous findings that developmental-stage neurons exhibit species-specific characteristics between primates and rodents [32].

Overall, these findings suggest that the latent space learned by MUSE captures both conserved features across species and species-specific characteristics associated with developmental stages.

### Visualization of Motif Activity by chromVAR

To assess whether the MUSE latent space preserves lineage-associated motif activity across species, we performed motif enrichment analysis using chromVAR [33] on multi-omics data from macaque and mouse. This approach enables the identification of lineage-specific regulatory modules and reveals patterns that are consistently observed across species.

We first examined whether differences in bHLH regulatory modules between excitatory neuronal lineages and ventral forebrain-derived lineages are reflected in the MUSE latent space. The *NEUROD* family comprises bHLH transcription factors that promote neuronal differentiation and maturation [34]. Although motif MA0461.2 is annotated as corresponding to *ATOH1* [35], it represents a canonical E-box sequence that can be bound by multiple bHLH factors [36], including proneural bHLH proteins such as *NEUROG* and *NEUROD* [36, 37]. Consistent with this, the enrichment score of MA0461.2 increased along the excitatory neuron lineage (Fig. 3A).

**Fig. 3.**
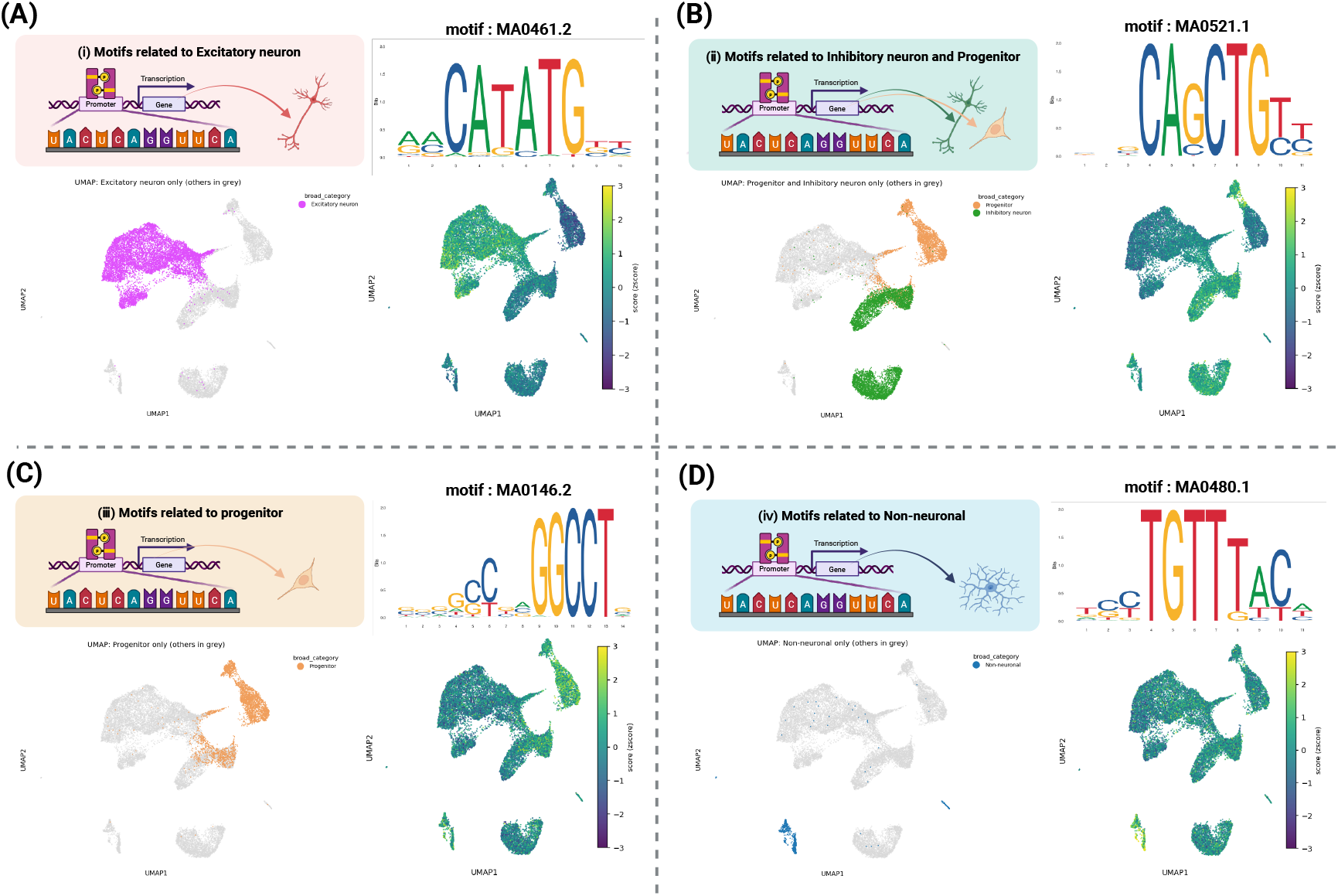
Motif enrichment analysis by chromVAR. Motif enrichment analysis using chromVAR performed on macaque and mouse multi-omics data projected onto the MUSE latent space. Enrichment scores of representative transcription factor motifs are shown across major lineages. (A) Motif MA0461.2 (E-box motif associated with proneural bHLH factors such as NEUROD) increases along the excitatory neuron (Ex) lineage. (B) Motif MA0521.1, bound by E-proteins cooperating with ASCL1, shows elevated activity in ventral forebrain-derived lineages including interneurons (IN) and oligodendrocyte lineage cells (OPC/OL). (C,D) Motif MA0146.2 (Zfx) is enriched in progenitor regions, whereas motif MA0480.1 (FOX family) is specifically enriched in non-neuronal populations. These patterns indicate that the MUSE latent space preserves conserved lineage-specific transcriptional regulatory architectures across species.

In contrast, E-proteins are class II bHLH transcription factors that form heterodimers with class II partners such as *ASCL1*, thereby enhancing DNA binding affinity and transcriptional activity [38]. E-proteins are known to bind motif MA0521.1 [35] and function as key regulators in ventral forebrain transcriptional programs. Moreover, the *ASCL1*–E-protein cooperative module has been reported to function not only in interneuron (IN) lineages but also in oligodendrocyte precursor cells (OPC) and oligodendrocytes (OL), contributing to bHLH-dependent programs during oligodendrocyte lineage specification and differentiation [39, 40]. Consistently, increased activity of motif MA0521.1 was observed in both IN and OPC/OL populations, consistent with ventral lineagespecific regulatory modules (Fig. 3B).

We further observed lineage-specific motif activity associated with progenitor and non-neuronal cell populations in the MUSE latent space. Motif MA0146.2 corresponds to *Zfx* [35], a transcription factor essential for maintaining selfrenewal capacity in progenitor cells [41]. In our analysis, MA0146.2 showed high enrichment scores in progenitor regions of the latent space, indicating active transcriptional programs associated with undifferentiated states (Fig. 3C).

In contrast, motif MA0480.1 corresponds to the *FOX* (Forkhead box) family [35], whose transcription factors are known to regulate homeostasis in non-neuronal cells such as endothelial and immune cells [42–44]. Accordingly, MA0480.1 exhibited increased activity specifically in non-neuronal regions of the latent space (Fig. 3D).

Taken together, these results indicate that the MUSE latent space recapitulates key transcriptional regulatory structures across lineages: a gradient of E-box activity centered on *NEUROD* in dorsal lineages, an *ASCL1*–*TCF12* cooperative module in ventral lineages, elevated *Zfx*-associated motif activity in progenitor regions, and increased *FOX* family motif activity in non-neuronal populations.

### Cross-Species Alignment Using MUSE Latent Space and SAMap

Using alignment scores computed in the shared latent space of human and mouse obtained from MUSE, we evaluated the validity of cross-species correspondence at a coarse-grained level, including progenitors, excitatory neurons, inhibitory neurons, and non-neuronal cells. Alignment scores were computed using the SAMap framework [45]. An overview of the workflow combining MUSE latent space and SAMap for alignment score computation is shown in Fig. S2. First, MUSE was used to learn a shared latent space that integrates RNA expression and ATAC accessibility into a unified coordinate system. Next, SAMap was applied to this latent space to estimate cell-cell correspondences as alignment scores (see Methods for details). Finally, cell types were grouped into four major categories (progenitor cells, excitatory neurons, inhibitory neurons, and non-neuronal cells), and cross-species consistency was evaluated within each category. This framework enables validation of cross-species alignment based on integrated representations that reflect both transcriptomic and epigenomic information.

We next present the alignment scores indicating the mapping of mouse cell types to human cell types (Fig. 4A). Overall, cross-species alignment was largely consistent within the same major categories (progenitors, excitatory neurons, inhibitory neurons, and non-neuronal cells). In particular, nonneuronal cells showed strong alignment between mouse and human, consistent with previous findings that non-neuronal cell types are more conserved across species [24].

**Fig. 4.**
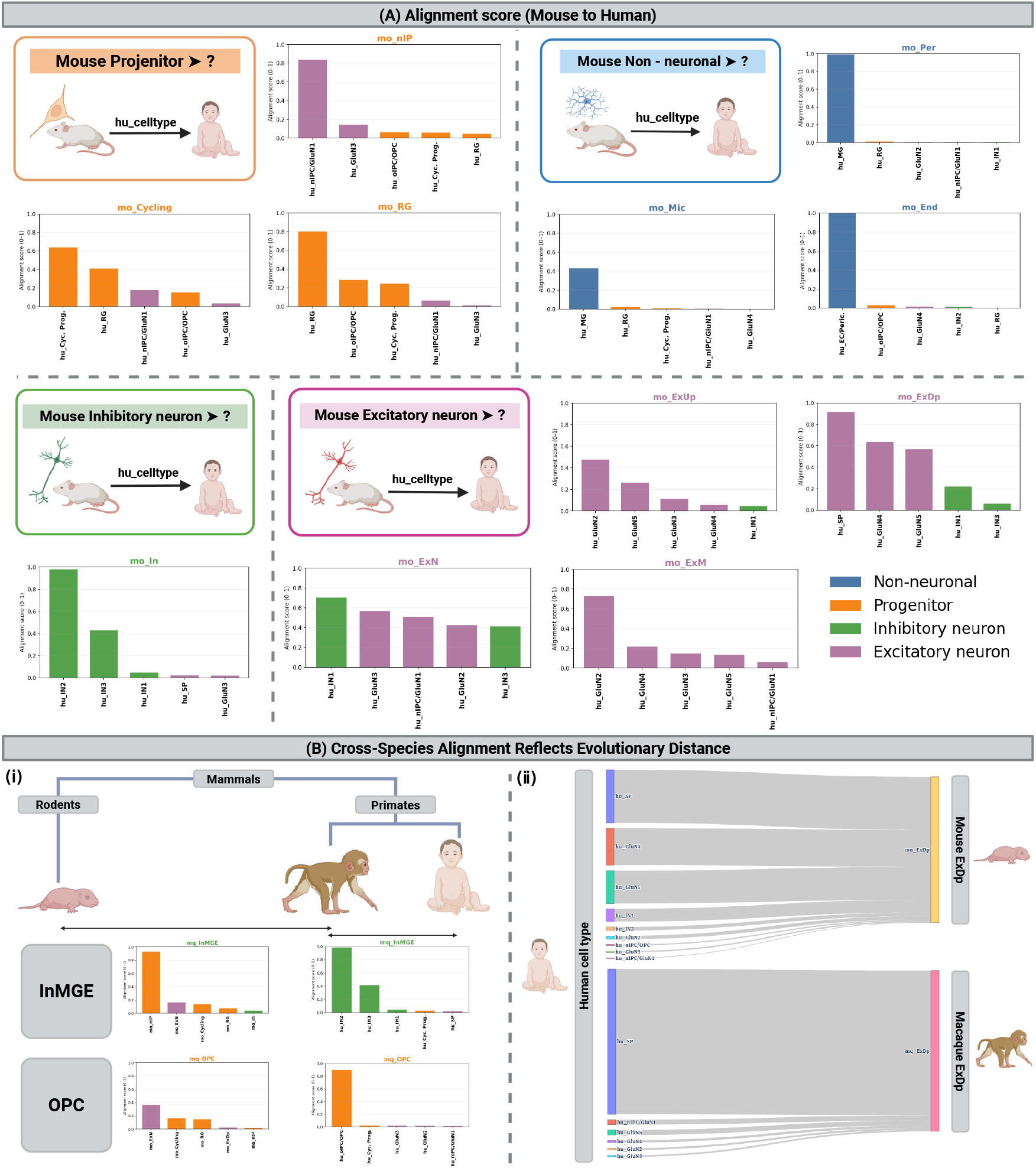
Estimation of cell-type alignment scores using SAMap. (A) Alignment scores between mouse and human cell types grouped into four major categories: progenitors, excitatory neurons (Ex), inhibitory neurons (IN), and non-neuronal cells. Most correspondences occur within the same broad category, with particularly strong conservation observed in non-neuronal cell types. (B) Effect of phylogenetic distance on cross-species alignment. (i) Alignment scores for macaque InMGE and OPC cells with human and mouse cell types. Closely related species (human–macaque) show dominant correspondence within the same lineage, whereas more distant comparisons (macaque–mouse) exhibit broader cross-lineage correspondences. (ii) Alignment of ExDp cells across species, showing near one-to-one correspondence between macaque and human SP cells, while mouse ExDp cells align with multiple human cell types.

However, cross-category alignments were also observed. Mouse excitatory neurons (ExN) showed high alignment scores with human inhibitory neurons (IN1), despite their functional differences. In progenitor populations, mouse nIP cells exhibited high alignment scores with human nIPC/GluN1 cells, which represent transitional neuronal states including early-stage neurons [46]. These correspondences are likely driven by proximity along developmental trajectories rather than strict cell-type equivalence.

Next, to assess whether alignment strength consistent with phylogenetic distance, we computed alignment scores using MUSE latent spaces for human–macaque and macaque– mouse comparisons, and evaluated the relationship between phylogenetic distance and cross-species alignment (Fig. 4B).

First, for macaque cell types such as InMGE and OPC, we computed alignment scores with corresponding human and mouse cell types (Fig. 4B(i)). In both cases, alignment with the evolutionarily closer human showed strong correspondence within the same lineage, whereas alignment with the more distant mouse exhibited cross-lineage correspondences.

We further visualized how macaque and mouse ExDp cells align to human cell types (Fig. 4B(ii)). Macaque ExDp cells showed strong alignment with human subplate neurons (SP), whereas mouse ExDp cells exhibited moderate alignment with multiple human cell types. These results are consistent with previous findings that closely related species tend to exhibit one-to-one cell-type correspondences, while more distantly related species show many-to-many relationships that reflect similarities at the level of regulatory modules, forming cell-type families [45].

### MUSE Enables Cross-Species Alignment Incorporating Regulatory Modules

We applied SCENIC+ to human and mouse datasets independently to identify active eRegulons in each species (Fig. 5). Next, we performed Gene Ontology (GO) analysis on the target genes of each eRegulon and assigned functional labels, enabling us to determine which functional modules are specifically active in each cell type (Fig. 5A).

**Fig. 5.**
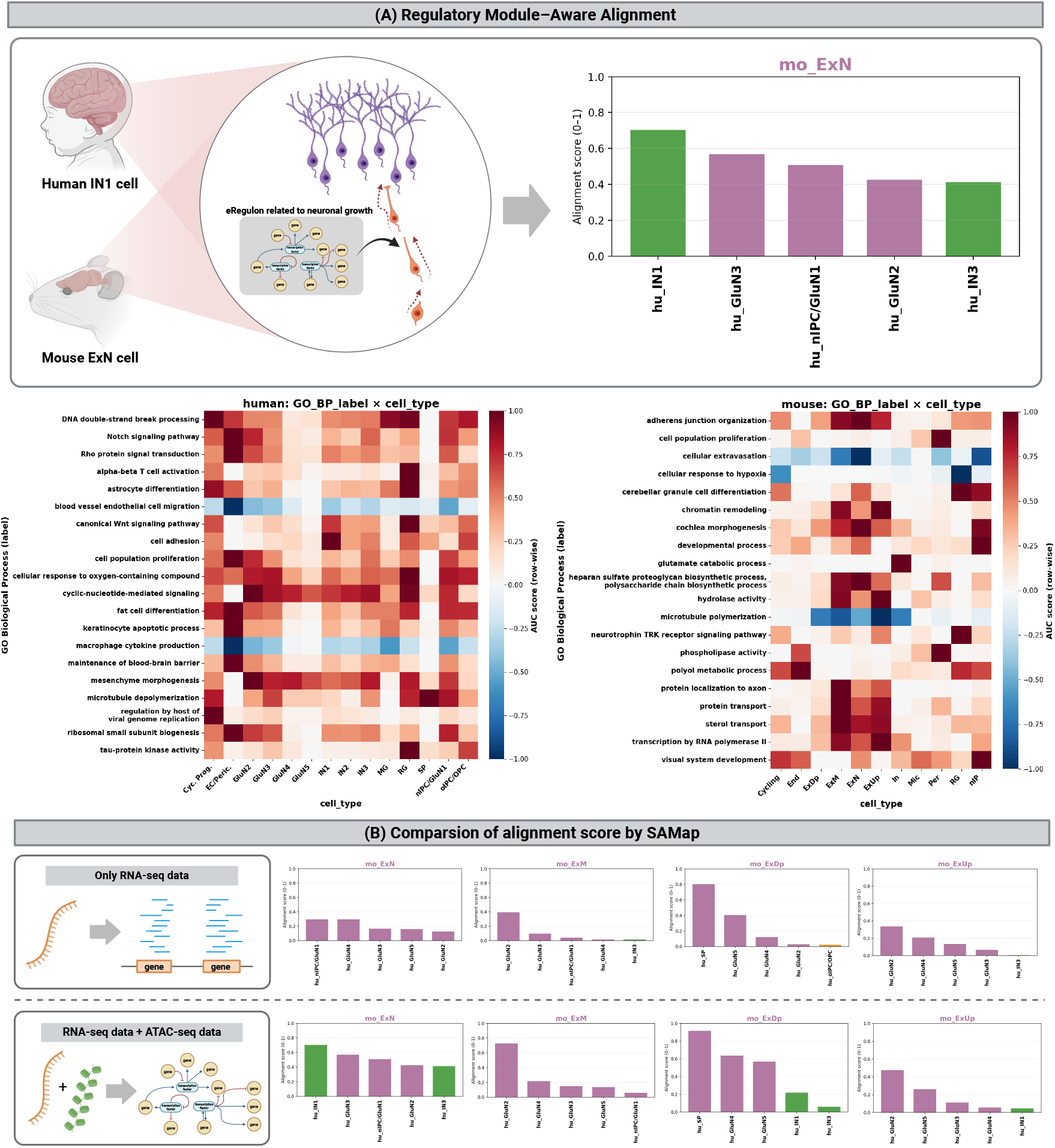
Cross-species alignment incorporating regulatory modules. A) Functional characterization of active eRegulons identified independently in human and mouse using SCENIC+. Gene Ontology (GO) enrichment analysis was performed on target genes of each eRegulon to assign functional labels, enabling comparison of regulatory modules active in specific cell types across species. Notably, regulatory modules related to cell adhesion, synapse formation, and neural circuit organization are active in both human inhibitory neurons (IN1) and mouse excitatory neurons (ExN). (B) Cross-species alignment obtained using SAMap based solely on RNA expression data. In contrast to the MUSE-based alignment (Fig. 4), RNA-based mapping predominantly aligns cell types within the same lineage and rarely detects cross-lineage correspondences such as IN1–ExN.

In cross-species alignment based on MUSE and SAMap, human inhibitory neurons (IN1) and mouse excitatory neurons (ExN) showed high alignment scores despite belonging to different lineages (Fig. 4A, Fig. 5A). To investigate this observation, we compared eRegulon activity between these two cell types based on their functional annotations. In human IN1 cells, eRegulons associated with functional modules such as the canonical Wnt signaling pathway, cell adhesion, and adherens junction organization showed relatively high activity (Fig. 5A). These modules are known to be involved in axon extension, synapse formation, dendritic spine maturation, and stabilization of excitatory synapses [47, 48]. In contrast, mouse ExN cells showed high activity in eRegulons associated with morphogenetic and extracellular matrix-related programs, including GO terms such as cochlea morphogenesis and heparan sulfate proteoglycan biosynthetic processes (Fig. 5A). These modules are involved in neural circuit formation and axon guidance through regulation of Wnt signaling, cell adhesion, cytoskeletal dynamics, and heparan sulfate chain structure [49–51].

Taken together, these results suggest that both IN1 and ExN exhibit convergence in functionally related regulatory programs associated with structural and morphogenetic aspects of neural circuit formation, including cell adhesion, synapse formation, and axon guidance. This suggests that, despite their inhibitory versus excitatory lineage differences, these cell types exhibit convergence at the level of functional regulatory modules. The high alignment scores between IN1 and ExN in the MUSE latent space can therefore be interpreted as reflecting such regulatory module-level similarity.

In contrast, when SAMap was applied using RNA data alone, cross-species alignment was largely restricted within the same lineage, and cross-lineage correspondences such as IN1–ExN were rarely detected (Fig. 5B). Previous studies have suggested that RNA-based mapping is strongly dominated by lineage signals and may fail to capture similarities in conserved regulatory modules even when they exist [52]. Our results are consistent with this observation.

Furthermore, prior work[24] has shown that as phylogenetic distance increases, the conservation of cell-type marker genes decreases, while comparisons based on regulatory module structure become more informative. In our study, the increased prevalence of cross-lineage alignments with greater phylogenetic distance suggests this notion. These findings suggest that the cross-species latent representations learned by MUSE, which integrate chromatin accessibility and transcriptomic information, enable the identification of conserved and divergent regulatory modules that cannot be captured by single-omics approaches alone.

## Discussions

In this study, we proposed MUSE, a method for integrating cross-species multi-omics data, and demonstrated its effectiveness using RNA and ATAC datasets derived from the developing cerebral cortex of human, macaque, and mouse. MUSE extends the GLUE framework [53], which was originally designed for single-species integration, by constructing a cross-species guidance graph. Specifically, it preserves gene regulatory networks constructed within each species while integrating cross-species correspondences based on amino acid sequence similarity and ortholog information. This design enables the suppression of species-specific batch effects while capturing cross-species latent structure consistent with conserved regulatory modules, as well as lineagespecific regulatory programs, within a shared latent space.

Benchmarking results suggest that the latent space learned by MUSE captures conserved transcriptional programs across species while simultaneously preserving species-specific transcriptional states associated with developmental stages. In particular, in ExDp cells, mouse cells that align with macaque ExDp cells showed high expression of genes related to synapse formation and circuit maturation, whereas mouse cells that do not align exhibited elevated expression of genes associated with developmental processes such as cortical layer formation and cell migration. These findings indicate that both conserved functional maturation programs (e.g., synapse formation and circuit maturation) and species-specific developmental programs are represented within the latent space.

Furthermore, motif enrichment analysis using chromVAR revealed that the MUSE latent space preserves biologically interpretable, lineage-associated motif activity patterns. These include a gradient of E-box activity associated with *NEUROD*-family bHLH factors in dorsal lineages and *ASCL1*–E-protein cooperative modules in ventral lineages. These results suggest that MUSE latent representations integrating chromatin accessibility and transcriptomic information simultaneously preserve both conserved and species-specific regulatory modules that govern developmental lineages. In this regard, MUSE provides a major advantage in its ability to capture both conserved regulatory modules and developmental-stage-dependent divergence across species.

Cross-species alignment based on MUSE and SAMap revealed that most cell types are aligned within the same lineage, while some cases exhibited moderate alignment across different lineages, such as between inhibitory neurons (IN1) and excitatory neurons (ExN). Further analysis showed that they share functionally related regulatory programs related to cell adhesion, axon guidance, and morphogenesis, suggesting that transient cellular states during development can lead to molecular similarities that are not fully explained by lineage differences. In contrast, cross-species alignment based on RNA data alone rarely detected such cross-lineage correspondences.

These findings demonstrate that MUSE latent representations enable the identification of state-dependent and regulatory module-level conservation that can be difficult to capture using single-omics approaches alone. Thus, MUSE enables the exploration of biological relationships beyond lineage-level correspondence.

Overall, our results show that MUSE enables (i) reduction of species-derived batch effects, (ii) extraction of species-specific features, and (iii) cross-species alignment that accounts for regulatory modules, thereby providing a comprehensive framework for cross-species multi-omics integration. Despite these strengths, several limitations of MUSE remain. First, the robustness of the proposed method with respect to species diversity has not been fully evaluated. In this study, real-data applications were limited to three species: human, macaque, and mouse. We did not assess the performance of the method on other species. Previous studies have suggested that integration performance may vary depending on the phylogenetic distance between species [54]. Therefore, it will be important to evaluate the robustness of MUSE in settings involving more distantly related species, such as combinations of primates and fish.

Second, the execution workflow of the proposed method remains relatively complex. In the current framework, feature graphs must be constructed separately for each species, and ortholog relationships as well as feature similarities must be computed for all pairs of species. While this does not impose a substantial burden in scenarios involving pairwise comparisons between human and model organisms, as in this study, the computational cost and workflow complexity increase substantially when integrating a larger number of species simultaneously. Future work should therefore focus on revising the current implementation, which relies on pairwise computations across all species, and developing a more scalable framework that mitigates the increase in complexity as the number of species grows.

## Methods

### MUSE framework

First, we describe the framework of MUSE, which extends the existing multi-omics data integration method GLUE (graph-linked unified embedding)[53] to cross-species multi-omics data integration. Let the omics data matrix of the *k* ∈ 𝒦-th modality for species *s* ∈ 𝒮 be denoted by 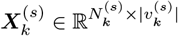. Here, 𝒮 denotes the index set of species, 𝒦 denotes the index set of omics modalities, 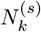 represents the number of cells of the *k*-th omics data for species 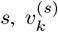 denotes the set of features, and 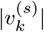 represents its cardinality. MUSE is a VAE (Variational AutoEncoder)[55]-based method that integrates heterogeneous multi-omics data 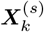 by learning a shared latent space.

In MUSE, encoders for embedding the data and decoders for reconstructing the data are defined for each species and each omics modality. The encoder and decoder for the *k*-th omics data matrix of species *s* are denoted as 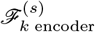 and 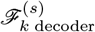, respectively. By inputting the data ma-trix 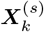 into the encoder, a low-dimensional representation 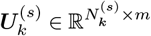 is obtained. Here, *m* denotes the dimensionality of the latent variables, typically set to *m* = 50.

The relationships across different modalities are captured by a guidance graph 𝒢^(*s*)^ constructed for each species *s*. This guidance graph is a prior knowledge-based graph in which nodes correspond to features from different omics modalities (e.g., genes and peaks), and edges and weights are assigned based on distances computed from feature coordinates[53]. In addition to capturing relationships between modalities, MUSE constructs a cross-species guidance graph 𝒢 by integrating the guidance graphs 𝒢 ^(*s*)^ of each species based on protein embedding similarity and ortholog information. Details are described in Section 5.2.

Using the embedding 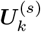 of the data matrix 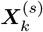 and the graph embedding 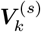 obtained by applying a GCN to the graph 𝒢, MUSE integrates cross-species multi-omics data by learning representations such that the data matrix can be re-constructed from their inner products.

The overall training of MUSE is performed based on the following objective function:

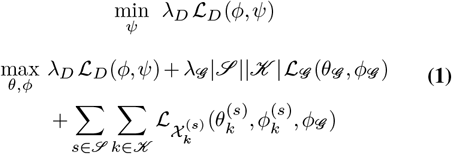

Here, 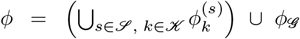 and 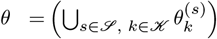 ∪ *θ*_*𝒢*_ denote the sets of parameters for the data encoders and graph encoder, and the data decoders and graph decoder, respectively. In MUSE, the parameters *ϕ* and *θ* are learned by maximizing the objective function Eq. (1). The objective function consists of three components: the adversarial loss ℒ_*D*_, the graph reconstruction loss ℒ_*G*_, and the data reconstruction loss 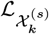.

The first term is the adversarial loss, defined as follows:

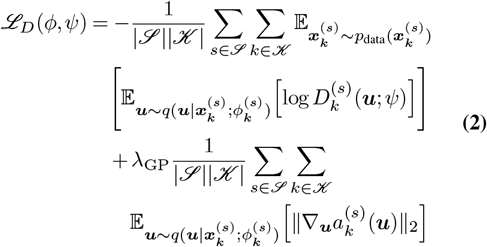

*ψ* denotes the parameters of the discriminator, and 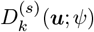 represents the probability that the data *x* corresponding to the latent variable ***u*** belongs to the *k*-th omics data of species 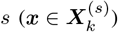. Moreover, *p*_data_ denotes the data distribution and *q* denotes the posterior distribution.

Adversarial learning is formulated as a min–max optimization problem, where the discriminator and generator are trained in a competitive manner. Since it is known that the training of the parameters *ψ* and *ϕ* becomes unstable under such a setting, we introduce a zero-centered gradient penalty to stabilize the training [56]. The second term in Eq. (2) corresponds to this gradient penalty, which suppresses excessive sensitivity of the discriminator in the latent space and improves the stability of adversarial learning.

The second term is the graph reconstruction loss, defined as follows:

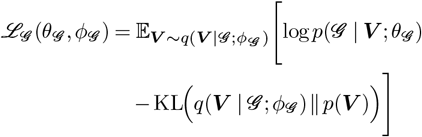

Here, 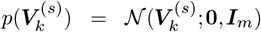 and 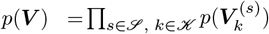 denote the prior distribution of the graph, and *q* denotes the posterior distribution. The model is trained such that the original graph structure can be reconstructed from the graph embeddings.

The third term represents the data reconstruction loss, defined as follows:

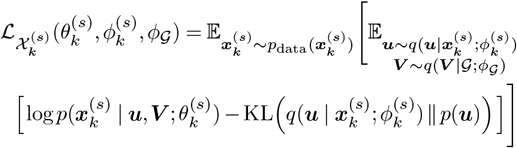

Here, *p*(***u***) = 𝒩(***u***; **0, *I***_*m*_) denotes the prior distribution of the latent variable ***u***. By integrating the omics data embeddings 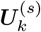 and graph embeddings 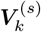 for each species, the model reconstructs the omics data.

### Cross Species guidance graph

The guidance graphs of individual species are integrated based on gene homology to construct a cross-species guidance graph. The structure of this cross-species guidance graph enables the characterization of gene homology across species from both continuous similarity and discrete correspondence perspectives. Specifically, similarity based on amino acid sequences is represented as a continuous measure, while orthologous relationships between genes are represented as discrete correspondences.

To represent amino acid sequence similarity, we employ the protein language model ESM2 [23], which takes amino acid sequences as input and outputs high-dimensional embeddings that capture information related to the threedimensional structure of proteins. By computing the similarity between vectors obtained from ESM2, we quantitatively evaluate gene similarity across species based on amino acid sequences.

Orthologous relationships between genes are identified by referring to ortholog information obtained from gene IDs.

The cross-species guidance graph is constructed by augmenting each species-specific guidance graph 𝒢 ^(*s*)^ with inter-species edges between gene nodes 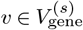 and 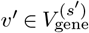. These edges include weighted edges based on ESM2 embedding similarity, denoted as 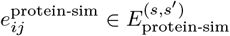, and weighted edges based on ortholog relationships, denoted as 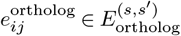.

We first describe the construction of weighted edges based on ESM2 embedding similarity. ESM2 is a Transformer-based protein language model that treats amino acids as tokens [23]. In this study, we use the pretrained model esm2_t6_8M_UR50D to compute sequence embeddings. We then define a similarity function 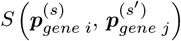 for computing similarity between vectors obtained from ESM2.

Here,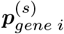 and 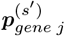 denote the protein vectors of gene *i* in species *s* and gene *j* in species *s*′, respectively, and the similarity function *S* is defined using the following RBF (Radial Basis Function) kernel:

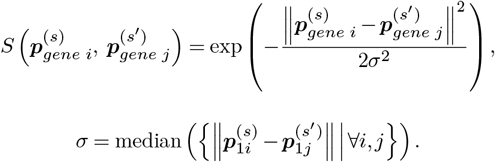

The hyperparameter *σ* in the RBF kernel is set to the median of the Euclidean distances between protein vectors to prevent the similarity distribution from becoming excessively skewed and to ensure that differences in similarity are adequately reflected. If the value of the similarity function exceeds a threshold *c*, a weighted edge is added between gene nodes across species. Thus, the set of weighted edges based on ESM2 embedding similarity is defined as:

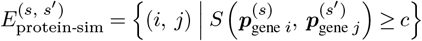

Next, we describe the assignment of weighted edges based on ortholog information. Ortholog genes are genes from different species that originate from a common ancestral gene. Ortholog information is extracted from a database (Ensembl Compara)[57] using gene IDs, and edges with weight 1 are added between gene nodes 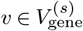 and 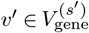 that are orthologous. If an edge has already been assigned based on ESM2 similarity, it is overwritten with weight 1. The set of such edges is defined as 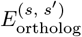.

Finally, the set of edges added between gene nodes across species is defined as the union:

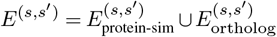

### Data processing

Data preprocessing and feature selection were performed using the existing tools Seurat and Link-Peaks [18, 58]. Variable genes (HVGs) were identified from the RNA assay using Seurat, and the top 3,000 genes were selected as the candidate set. Next, the analysis was restricted to protein-coding genes, and sequence embeddings for each gene were computed using a pre-trained protein language model, ESM2 (esm2_t6_8M_UR50D). Previous studies have shown that sequence representations obtained through selfsupervised learning of protein language models capture biologically meaningful information such as secondary and tertiary protein structures [59].

After assigning gene embeddings, genes with assigned embeddings were compiled into a list and used as the input gene set for MUSE (human: 2333 genes, macaque: 1852 genes, mouse: 2141 genes). For ATAC data, peak–gene associations between the selected genes and all peaks were inferred using LinkPeaks. For each peak–gene pair, a one-sided *z*-test was performed to assess whether the observed correlation was significantly greater than the background peak correlation distribution, and peaks with *p*-values less than 0.05 were retained. Furthermore, highly variable peaks were identified, and the union of statistically significant peaks and highly variable peaks was used as the input gene set for MUSE (human: 23109 peaks, macaque: 25087 peaks, mouse: 20985 peaks).

Based on these selections, AnnData objects were constructed: for RNA, including only genes with valid embeddings, and for ATAC, including only the union of significant peaks and highly variable peaks. Applying UMAP to the processed feature space showed that known cell-type labels were well separated in the embedding space, confirming that the preprocessing preserved biological signals appropriately (Figure **??**).

### Segmentation of Cell Types

The pre-annotated cell-type labels were grouped into broader categories as follows (Table 3.2).

RG are progenitor cells [60], and oRG represent the major progenitors of the expanded OSVZ in primates [61]. IPCs (nIP/gIP), which express EOMES/TBR2, function as intermediate neural progenitors and constitute the core lineage from RG (PAX6) to excitatory neurons (TBR1) [62]. OPC/oIPC are progenitors of the oligodendrocyte lineage, and in the human fetal cortex, EGFR^+^ pre-OPCs have been shown to emerge along the oRG continuum [63]. Based on these findings, the following cell types were classified as progenitor cells: in human, oIPC/OPC, RG, and Cyc.Prog; in macaque, Cycling, RG, nIP, gIP, and OPC; and in mouse, Cycling, RG, and nIP.

Endothelial cells (End) and pericytes (Per) are mural cells derived from mesodermal or ectodermal origins and do not belong to the neuroepithelial lineage [64][65]. Microglia originate from yolk sac-derived macrophage lineages during early embryogenesis and are phylogenetically independent of the neuroepithelium [66]. VLMC (vascular leptomeningeal cells) are fibroblast-like cells located in perivascular and meningeal regions, reflecting a mesenchymal origin distinct from the nervous system [67]. Based on these characteristics, the following cell types were classified as non-neuronal cells: in human, MG and EC/Peric.; in macaque, End, Per, Mic, and VLMC; and in mouse, End, Per, and Mic. These cell populations are known to be clearly separated from neuronal lineages at the transcriptomic level and form distinct clusters [68].

Ex and GluN represent excitatory neurons undergoing differentiation during embryonic development, and SP (subplate) corresponds to an early-born population of predominantly glutamatergic neurons that contribute to circuit relay and wiring control [69]. Accordingly, the following cell types were classified as excitatory neurons: in human, nIPC/GluN1, GluN2–GluN5, and SP; and in both macaque and mouse, ExDp, ExM, ExN, and ExUp.

InMGE, InCGE, and IN1–3 were treated as inhibitory neuron lineages. These cells originate from the MGE/CGE of the ventral telencephalon and are supplied to the cortex via tangential migration, as established in both classical and recent studies [70][71]. Therefore, the following cell types were classified as inhibitory neurons: in human, IN1–IN3; in macaque, InCGE and InMGE; and in mouse, In.

### Evaluation Metrics

As evaluation metrics, we used BatchScore and BioScore from scIB. BatchScore and BioScore consist of multiple metrics and are computed by scaling each metric to the range [0,1] and averaging them, following the procedure described in [72]. Each evaluation was repeated 20 times using random subsampling of 80% of the cells, and the final score was obtained as the average over repetitions[72]. BatchScore is a comprehensive metric that quantifies the extent to which species (batch) information is removed and how well cells from different species are mixed in the embedding space. A higher value indicates a smaller influence of species differences. In contrast, BioScore is a comprehensive metric that evaluates how well cell-type structure is preserved. A higher value indicates that biologically meaningful cell-type separation is well maintained.

The definitions of the individual metrics constituting BatchScore and BioScore are given below.

Let the embedding obtained from MUSE be ***Z*** ∈ ℝ^*n×m*^, and let *𝓏*_*i*_ denote the embedding of cell *i*. Let *b*_*i*_ ∈ {1, …, *B*} be the species label and *y*_*i*_ ∈ {1, …, *C*} be the cell-type label. Let 𝒩_*k*_(*i*) denote the *k*-nearest neighbors and *d*(*·,·*) denote a distance function. In this study, we use the Euclidean distance for *d*(*·,·*). Let *ĉ*_*i*_ denote cluster assignments, and when necessary, we use the top *d* principal components *H* = [*h*^(1)^, …, *h*^(*d*)^] obtained by PCA.

We define a generalized label 𝓁_*i*_ ∈ {*b*_*i*_, *y*_*i*_}. When 𝓁_*i*_ = *b*_*i*_, the evaluation is based on species labels (iLISI, BatchASW, BatchARI, BatchNMI), and when 𝓁_*i*_ = *y*_*i*_, it is based on celltype labels (cLISI, CellTypeASW, CellTypeARI, CellTypeNMI). For a label 𝓁, let *C*_𝓁_ = {*i* |𝓁_*i*_ = 𝓁} denote the set of cells with label *𝓁*.

### iLISI/cLISI

A metric that evaluates local diversity of labels. For each cell *i*, the empirical distribution of label 𝓁 in its *k*-nearest neighbors is

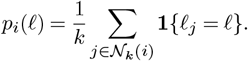

The inverse Simpson index is defined as

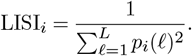

We aggregate this by taking the median across cells and apply the following scaling:

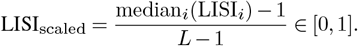

The final score is defined as

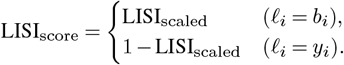

### kBET

A metric that evaluates whether the local batch composition matches the global batch composition using a statistical test. Let 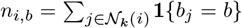 be the count of species labels in the neighborhood of cell *i*, and let 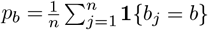 be the global proportion. The expected count is *E*_*i,b*_ = *kp*_*b*_, and the chi-square statistic is

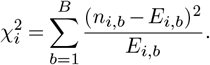

**Table 1.**
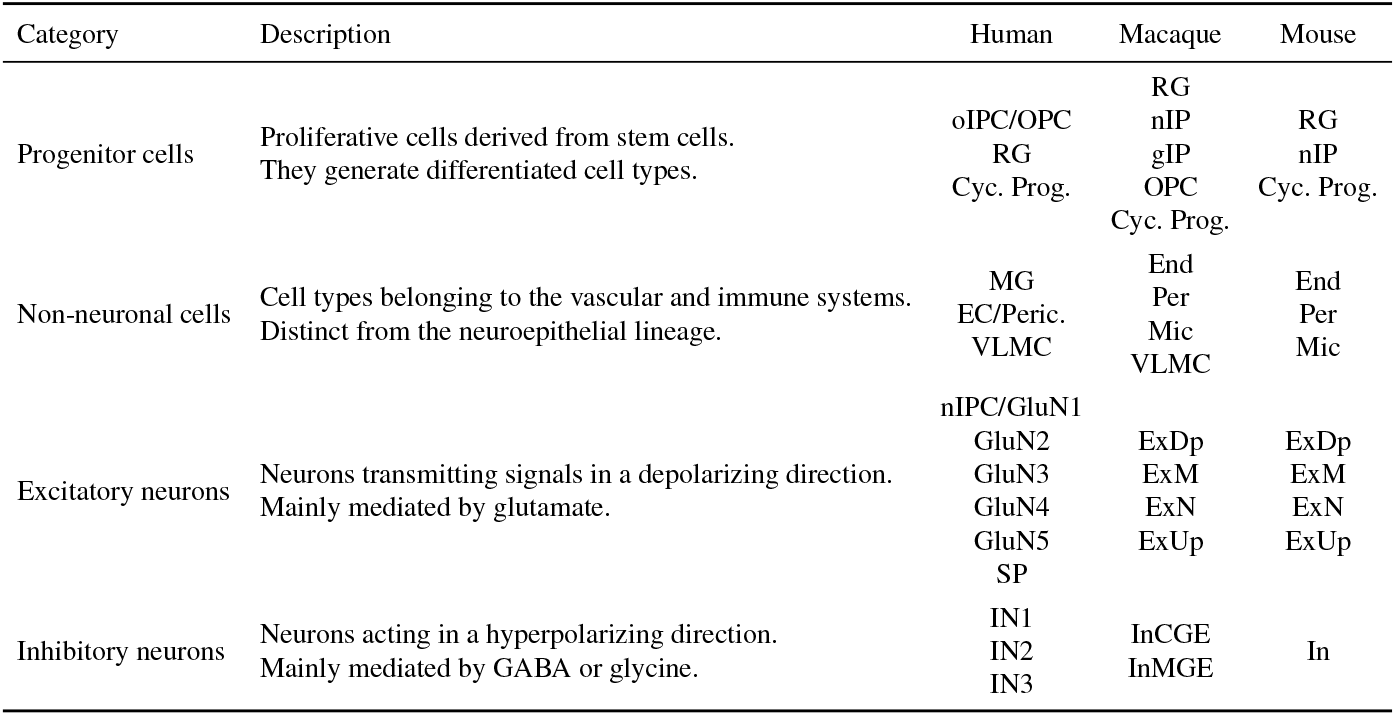
Major cell-type categories and their correspondence across species.

| Category | Description | Human | Macaque | Mouse |
| --- | --- | --- | --- | --- |
| Progenitor cells | Proliferative cells derived from stem cells.<br>They generate differentiated cell types. | oIPC/OPC<br>RG<br>Cyc. Prog. | RG<br>nIP<br>gIP<br>OPC<br>Cyc. Prog. | RG<br>nIP<br>Cyc. Prog. |
| Non-neuronal cells | Cell types belonging to the vascular and immune systems.<br>Distinct from the neuroepithelial lineage. | MG<br>EC/Peric.<br>VLMC | End<br>Per<br>Mic<br>VLMC | End<br>Per<br>Mic |
| Excitatory neurons | Neurons transmitting signals in a depolarizing direction.<br>Mainly mediated by glutamate. | nIPC/GluN1<br>GluN2<br>GluN3<br>GluN4<br>GluN5<br>SP | ExDp<br>ExM<br>ExN<br>ExUp | ExDp<br>ExM<br>ExN<br>ExUp |
| Inhibitory neurons | Neurons acting in a hyperpolarizing direction.<br>Mainly mediated by GABA or glycine. | IN1<br>IN2<br>IN3 | InCGE<br>InMGE | In |

We test whether the local distribution matches the global distribution. The acceptance indicator is 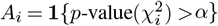, and the score is

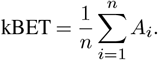

### BatchASW/CellTypeASW

A metric that evaluates cluster separation based on labels using silhouette width.

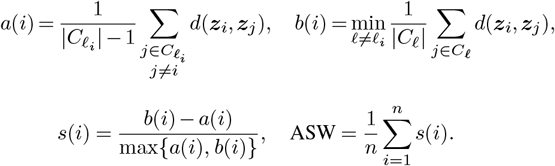

We normalize it as ASW_norm_ = (ASW + 1)*/*2 and define

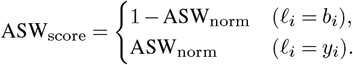

### BatchARI/CellTypeARI

Let *S*_*c*_ = {*i*: *𝓁*_*i*_ = *c*} be the set of cells with label *c*. We apply *k*-means clustering with the same number of clusters as label categories to embeddings in *S*_*c*_, and denote the assignments by 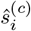. From the contingency table (*n*_*uv*_), the ARI is

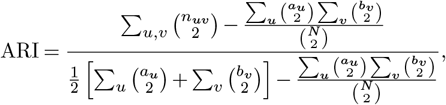

where 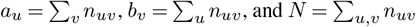. We average over labels:

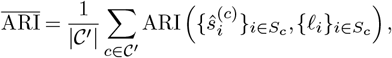

and define

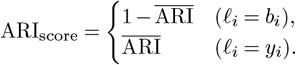

### BatchNMI/CellTypeNMI

A metric that evaluates agreement between clustering *ĉ* and labels *𝓁* using normalized mutual information:

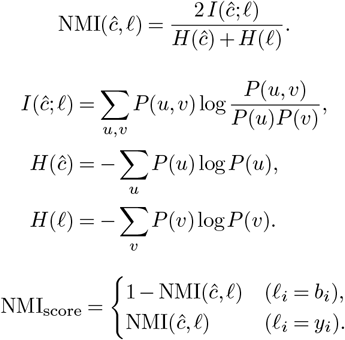

### GraphConn

A metric that evaluates how well cell-type structure is preserved based on the *k*NN graph. From the graph *G*, we take the induced subgraph *G*[*S*_*c*_] for each cell type *c*, which contains only nodes in *S*_*c*_ and the edges between them. Let LCC(*G*[*S*_*c*_]) be the largest connected component. Then

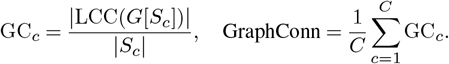

### PCR (Principal Component Regression)

A metric that evaluates how much species labels explain the principal components. We project ***Z*** onto *d* principal components and de-fine weights 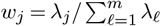. Using the one-hot matrix ***B*** as predictors:

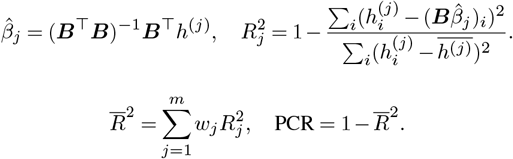

### Computation of Alignment Scores Using SAMap

Based on the SAMap algorithm proposed by Tarashansky et al. [45], we computed alignment scores between cell types by substituting the latent representations obtained from CS-GLUE into the alignment component of SAMap. Below, we present the formulation for a pair of two species (*i* ∈ {1, 2}).

#### Cell Representations and kNN Graphs

Let *n*_*i*_ denote the number of cells in species *i*, and let the representation vector of cell *u* be

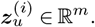

Here, *m* denotes the dimensionality of the representation vector, and in this study, we use the latent variables learned by CS-GLUE, i.e., 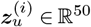.

We use cosine similarity as the measure of similarity between cells:

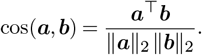

Based on this similarity, we define the top-*k* nearest neighbors. For a cell *u* in species *i*, let

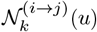

denote the set of top-*k* nearest neighbors in species *j* (intraspecies when *j* = *i*, inter-species when *j ≠ i*).

The adjacency matrix of the weighted *k*NN graph from species *i* to species *j*, 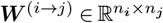, is defined as

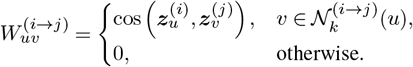

By normalizing each row to sum to 1, we define the intraspecies transition probability matrix 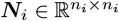 and the inter-species transition probability matrix 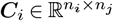 as

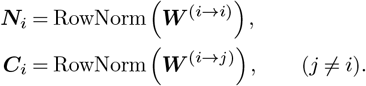

Here, RowNorm(*·*) denotes row-wise normalization such that each row sums to 1.

#### Construction of Mutual Nearest Neighbors (MNN) via k-hop Diffusion

In SAMap, to stabilize cross-species correspondence, the neighborhood structure is first smoothed within each species. Let ***N***_*i*_ be the intra-species transition probability matrix. Then, the *t*_*i*_-step diffusion is given by 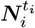, which represents the probability of reaching other cells after *t*_*i*_ steps of a random walk on the intra-species graph.

Next, each species-specific graph is clustered using the Leiden algorithm. Let *l*_*a*_ denote the size of the cluster *a* to which cell *u* belongs, and let 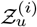 denote the neighborhood set of cell *u* on the graph. For each row of 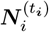, we retain only the *l*_*a*_ cells within 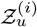, thereby constructing a smoothed neigh-borhood matrix 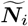 restricted to the local cluster.

Next, the transition probability matrix ***C***_*i*_ from species *i* to species *j* is aggregated using the local structure of species *j*. Specifically, the extent to which cell *u* in species *i* is connected to the neighborhood of cell𝓋 in species *j* is defined as

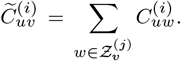

Finally, we compute the geometric mean of the aggregated matrices in both directions (1 *→* 2 and 2 *→* 1):

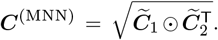

Here, ⨀ denotes the Hadamard (element-wise) product.

By retaining only the top-*k* edges for each row, we construct the final inter-species *k*NN graphs:

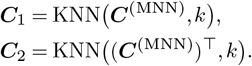

#### Cell-type Alignment Scores

Using the inter-species *k*NN graph 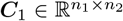, we define the directional alignment score 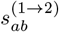 from cell type *a* in species 1 to cell type *b* in species 2. This score enables evaluation of cross-species correspondence at the level of major cell types such as progenitor, excitatory (Ex), inhibitory (IN), and non-neuronal cells. Let 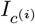 denote the set of cells belonging to cell type *c* in species *i*, and let 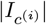 denote its cardinality. Then,

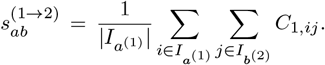

This score represents the extent to which cells of type *a* in species 1 are connected to cells of type *b* in species 2. A value close to 0 indicates weak correspondence, whereas a larger value indicates stronger alignment from species 1 to species 2.

### Implementation Details

In this study, cross-species multi-omics data were integrated using the `SCGLUE` function implemented in the `csglue` package. During model training, the following weights were assigned to the loss function: data reconstruction term *λ*_data_ = 1.0, KL regularization term *λ*_KL_ = 1.0, guidance graph regularization term *λ*_graph_ = 0.02, adversarial alignment term *λ*_align_ = 0.05, and gradient penalty term *λ*_GP_ = 0.1.

Optimization was performed using Adam, with an initial learning rate set to 2×10^*−*3^.

The mini-batch size during training was set to 128, and the ratio of training to validation data was set to 10:1. The maximum number of epochs, early stopping patience, and learning rate scheduling were automatically determined based on the dataset size using the default configuration of SCGLUE, and training was conducted until the early stopping criterion was satisfied.

All computational experiments in this study were conducted on a workstation equipped with Ubuntu 22.04 LTS and GPU acceleration.

## Competing interests

No competing interest is declared.

## Author contributions statement

Fuka Nakae(Data curation [Lead], Formal analysis [Lead], Methodology [Lead], Validation [Lead], Visualization [Lead], Writing—original draft [Lead], Writing—review & editing [Lead]), Shintaro Yuki(Formal analysis [Lead], Methodology [Lead], Validation [Supporting], Visualization [Supporting], Writing—original draft [Supporting], Writing—review & editing [Lead]), Zhenan Liu(Data curation [Supporting], Formal analysis [Lead], Methodology [Lead], Validation [Supporting], Visualization [Supporting], Writing—review & editing [Lead]), Chikara Mizukoshi(Data curation [Lead], Validation [Supporting], Visualization [Supporting], Writing—original draft [Supporting], Writing—review & editing [Supporting]), Shuto Hayashi(Formal analysis [Lead], Methodology [Lead], Writing—review & editing [Supporting]) Teppei Shimamura and Hiroshi Yadohisa (Project administration [Lead], Supervision [Lead], Writing—review & editing [Lead]).

## Data and Code Availability

In this study, experiments were conducted using processed publicly available datasets. Specifically, the human fetal cortex data were obtained from GEO accession GSE162170 (https://www.ncbi.nlm.nih.gov/geo/query/acc.cgi?acc=GSE162170), and the macaque and mouse data were obtained from GEO accession GSE241429 (https://www.ncbi.nlm.nih.gov/geo/query/acc.cgi?acc=GSE241429). The source code is currently being prepared and will be released in a public GitHub repository upon publication.

## Supplement

Supplementary Figures S1–S4 provide additional details of the analyses associated with the results presented in the main text. Figure S1 the overview of the baseline methods used for benchmarking, highlighting the framework of MUSE, which integrates species-specific guidance graphs by introducing cross-species edges based on ortholog relationships and similarity between protein sequence representations derived from ESM2. Figure S2 presents the analysis framework combining MUSE and SAMap, visualizing the procedure for identifying cell–cell correspondences in the shared latent space and computing alignment scores between cell types. Figure S3 shows the threshold selection procedure for ESM2 similarity in constructing the cross-species guidance graph. Figure S4 shows the data preprocessing pipeline, including feature selection for RNA and ATAC data, peak–gene association inference, and the preservation of cell-type structure as confirmed by UMAP visualization.

**Fig. S1.**
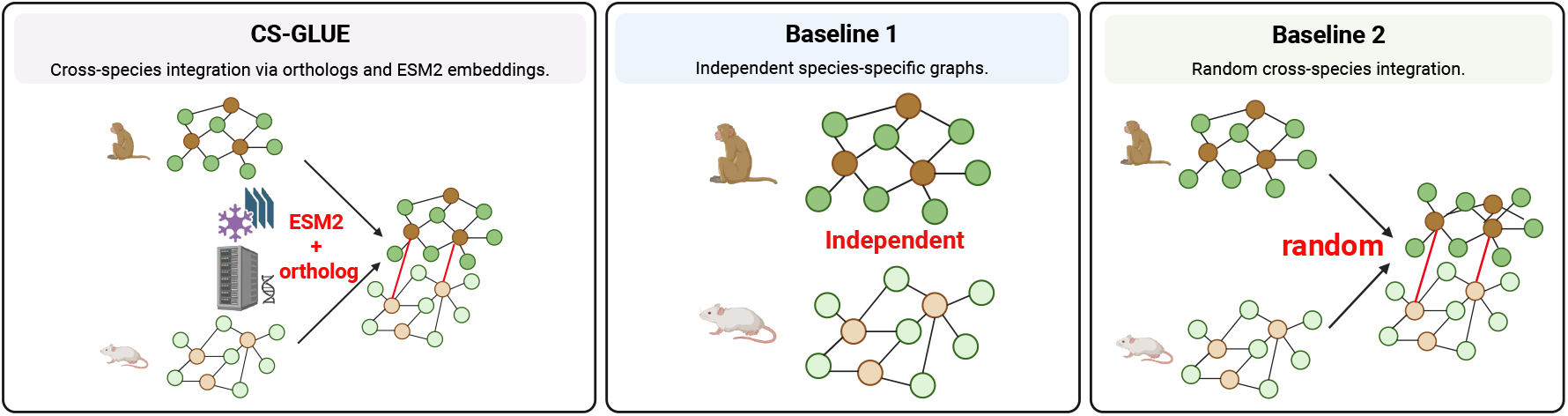
Baseline methods. MUSE integrates species-specific guidance graphs using cross-species edges defined by ortholog relationships and similarity between protein sequence representations. Baseline 1 uses independent species-specific graphs, whereas Baseline 2 introduces randomly assigned cross-species edges.

**Fig. S2.**
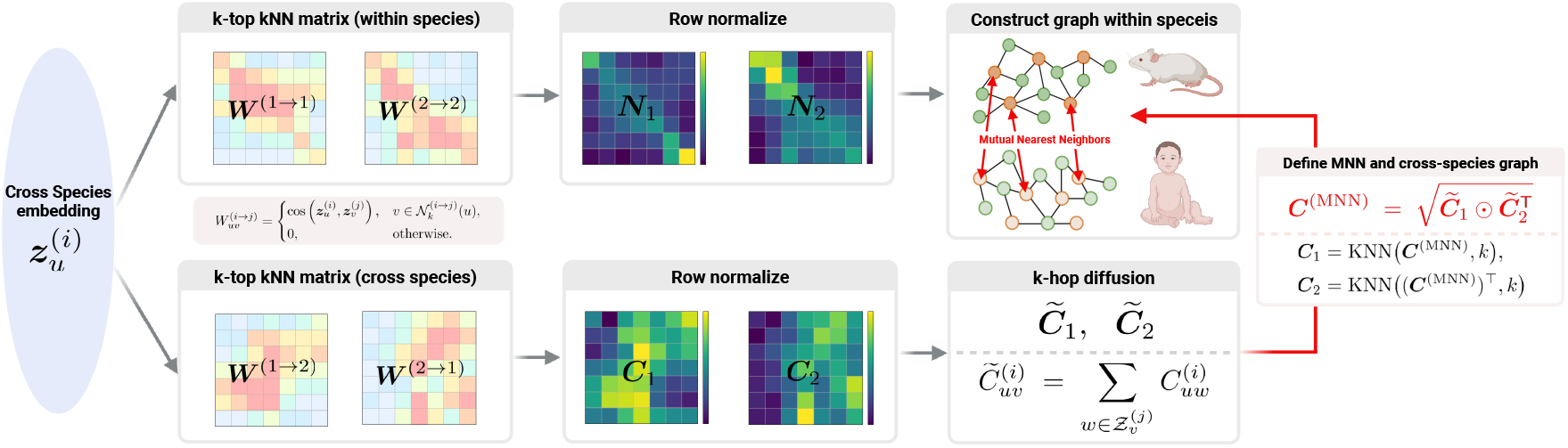
MUSE+SAMap framework. Workflow for computing cross-species alignment scores. MUSE first learns a shared latent space integrating RNA expression and ATAC accessibility across species. SAMap is then applied to this latent space to estimate cell–cell correspondences, which are summarized as alignment scores between cell types.

**Fig. S3.**
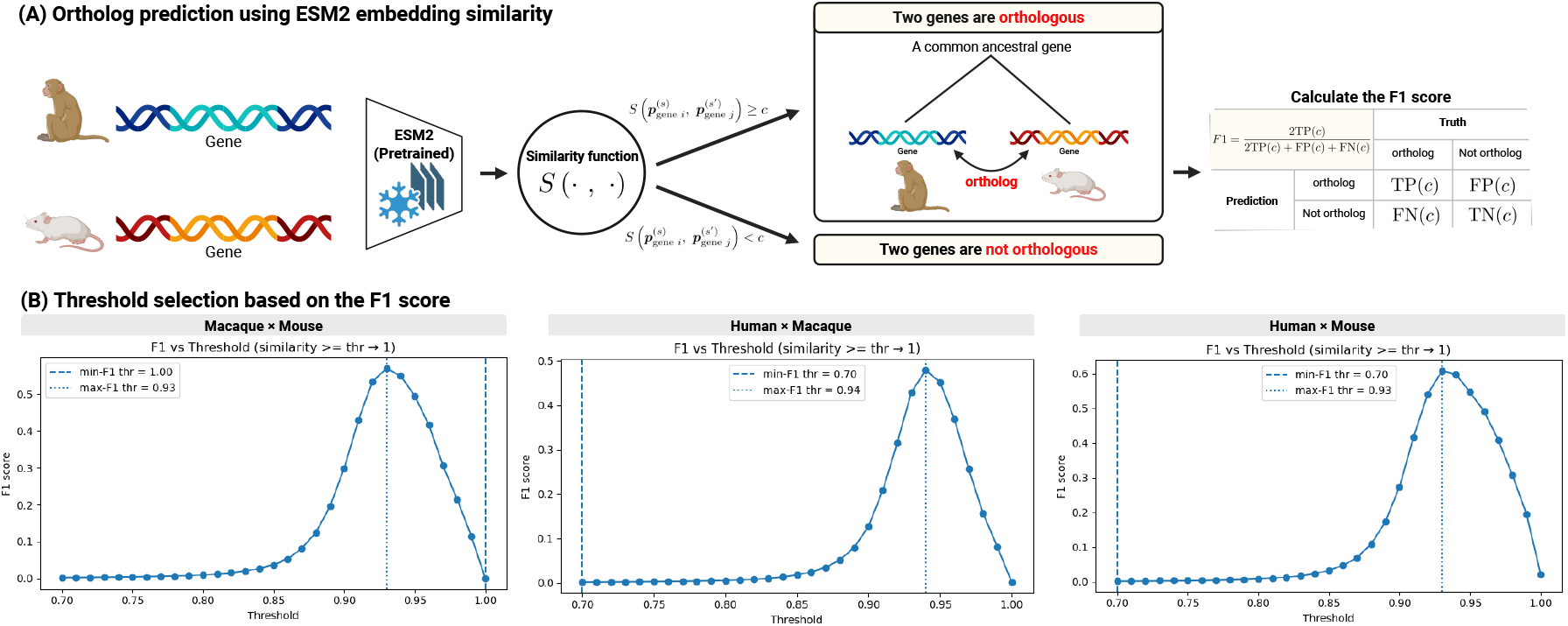
Threshold optimization. Using the computed similarities between ESM2 embeddings, we evaluated binary classification of whether each gene pair was orthologous across a range of thresholds from 0.70 to 1.00 in increments of 0.01. The threshold that achieved the highest F1 score was then selected as the cutoff for adding edges based on ESM2 embedding similarity.

**Fig. S4.**
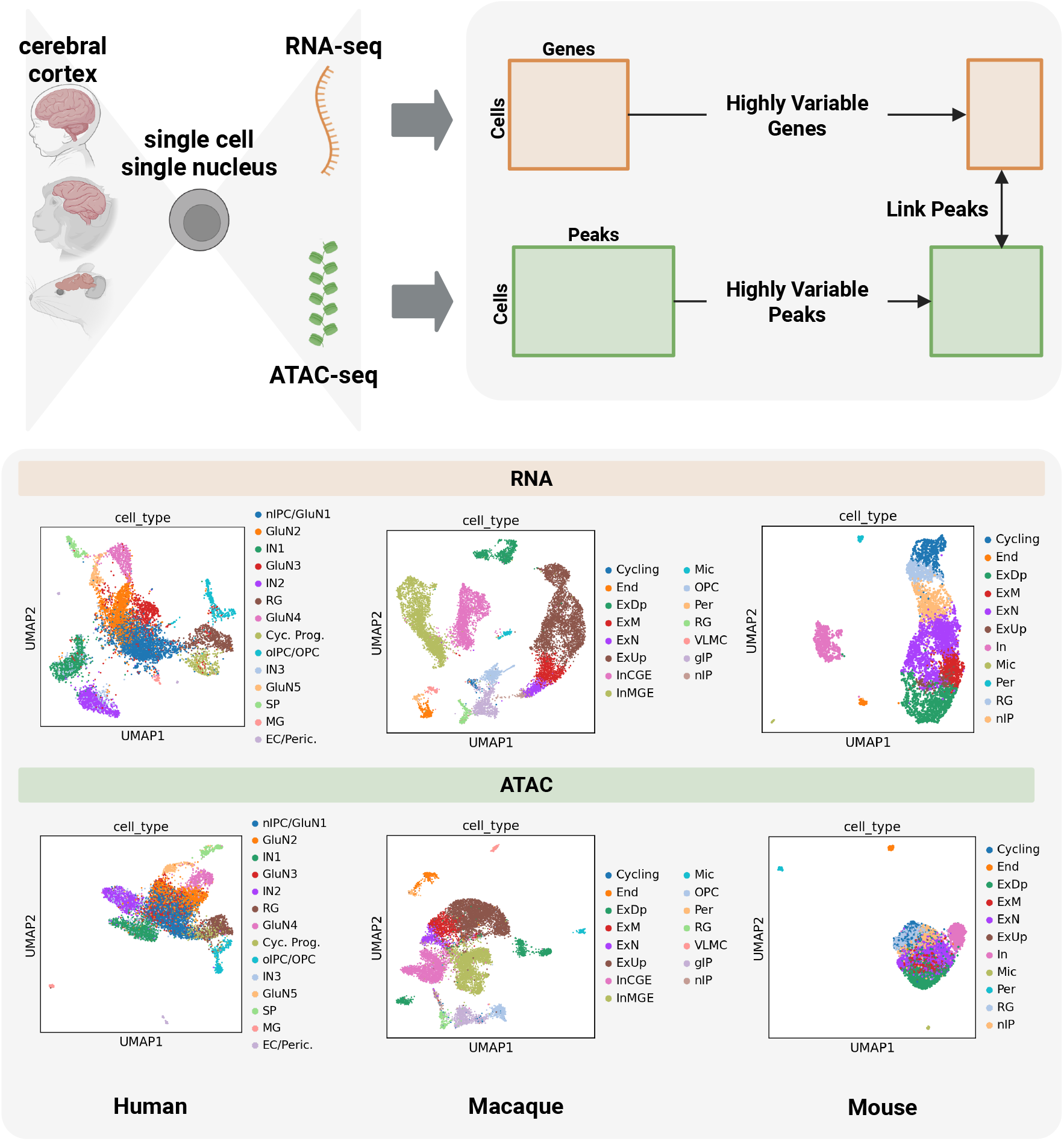
Data preprocessing pipeline. Single-cell RNA-seq and ATAC-seq data were processed using Seurat and LinkPeaks. For RNA, highly variable genes were selected and filtered to protein-coding genes with valid ESM2 embeddings. For ATAC, peak–gene associations were inferred, and significant peaks (p < 0.05) were combined with highly variable peaks. UMAP visualization shows clear separation of cell types across human, macaque, and mouse, indicating that biologically meaningful cell type structure is preserved in the integrated latent space.

## Notes

### Competing Interest Statement

The authors have declared no competing interest.

